# Comprehensive Approach to Simulating Large Scale Conformational Changes in Biological Systems Utilizing a Path Collective Variable and New Barrier Restraint

**DOI:** 10.1101/2023.03.26.534298

**Authors:** István Kolossváry, Woody Sherman

## Abstract

Conformational sampling of complex biomolecules is an emerging frontier in drug discovery. Advances in lab-based structural biology and related computational approaches like AlphaFold have made great strides in obtaining static protein structures for biologically relevant targets. However, biology is in constant motion and many important biological processes rely on conformationally-driven events. Conventional molecular dynamics (MD) simulations run on standard hardware, are impractical for many drug design projects, where conformationally-driven biological events can take microseconds to milliseconds or longer. An alternative approach is to focus the search on a limited region of conformational space defined by a putative reaction coordinate (i.e. path collective variable). The search space is typically limited by applying restraints, which can be guided by insights about the underlying biological process of interest. The challenge is striking a balance between the degree to which the system is constrained while still allowing for natural motions along the path. A plethora of restraints exist to limit the size of conformational search space, although each has drawbacks when simulating complex biological motions. In this work, we present a three-stage procedure to construct realistic path collective variables (PCV), and introduce a new kind of barrier restraint that is particularly well suited for complex conformationally-driven biological events, such as allosteric modulations and conformational signalling. The PCV presented here is all-atom (as opposed to C-alpha or backbone only) and is derived from all-atom MD trajectory frames. The new restraint relies on a barrier function (specifically, the scaled reciprocal function), which we show is particularly beneficial in the context of molecular dynamics, where near-hard-wall restraints are needed with zero tolerance to restraint violation. We have implemented our PCV and barrier restraint within a hybrid sampling framework that combines well-tempered meta-dynamics and extended-Lagrangian adaptive biasing force (meta-eABF). We use three particular examples of high pharmaceutical interest to demonstrate the value of this approach: (1) sampling the distance from ubiquitin to a protein of interest within the supramolecular Cullin-RING ligase complex, (2) stabilizing the wild-type conformation of the oncogenic mutant JAK2-V617F pseudokinase domain, and (3) inducing an activated state of the stimulator of interferon genes (STING) protein observed upon ligand binding. For (2) and (3), we present statistical analysis of meta-eABF free energy estimates and for each case, code for reproducing this work.

## 1. Introduction

Utilization and impact of molecular dynamics (MD) simulations in drug discovery continues to grow due to the insights that can be gained from understanding the motion of biomolecules at atomic resolution.^1–3^ However, traditional unbiased MD simulations are often insufficient to achieve the requisite level of sampling and accuracy fast enough to make an impact in drug discovery projects. As such, the use of enhanced sampling methods coupled with restraints in MD has become commonplace to improve conformational sampling and convergence of the free energy surface (FES). Restraining atomic positions or interatomic distances represents the most frequently used restraints, but many biologically relevant motions involve complex atomic transformations that cannot be treated effectively with such simple restraints and therefore require more sophisticated collective variables (CV) utilized in advanced sampling methods,^4,5^ which often require more sophisticated restraints.

One of the primary challenges faced when studying complex conformational transitions in proteins is elucidating a CV (e.g. some combination of distances, angles, torsions, contacts, accessible surface area, etc.) that captures the full complexity of the geometric perturbation. A particularly useful class of CVs, termed path CV (PCV), provides a flexible approach to model such transitions. Most PCVs employ a reduced representation model, such as a C-alpha (CA) trace, to generate a minimum potential energy path (MPEP), as introduced by Elber and Karplus. ^6–9^ The objective (and challenge) of PCV elucidation is to guide an all-atom simulation based on the MPEP in order to obtain a free energy surface (FES) in the region around the MPEP. In other words, the simulation evolves subject to certain restraints to keep it close to the MPEP while still sampling enough of the conformational space to obtain free energy estimates in the regions proximate to the MPEP.

The PCV often consists of an ordered set of equally spaced intermediate conformations called nodes. We extensively experimented with a number of different normal modes based PCVs and developed our own MPEP algorithm that extends Roux’s anisotropic network model (ANM).^10^ Details of our path algorithm are provided in Section 2, but it is instructive to note here that the main difference as shown on Figure 1 is that the ANM model presented in this work is smooth and differentiable everywhere, whereas the aforementioned path is based on a dual ANM model represented by two independent harmonic energy wells (one for each endpoint conformation) separated by a non-differentiable border where the two models intersect (see white border in the lower part of Figure 1. Our smooth-ANM (SANM) path is based on a unified ANM model, which features a genuine saddle point connecting the two energy wells as shown in the upper part of Figure 1.

**Figure 1:**
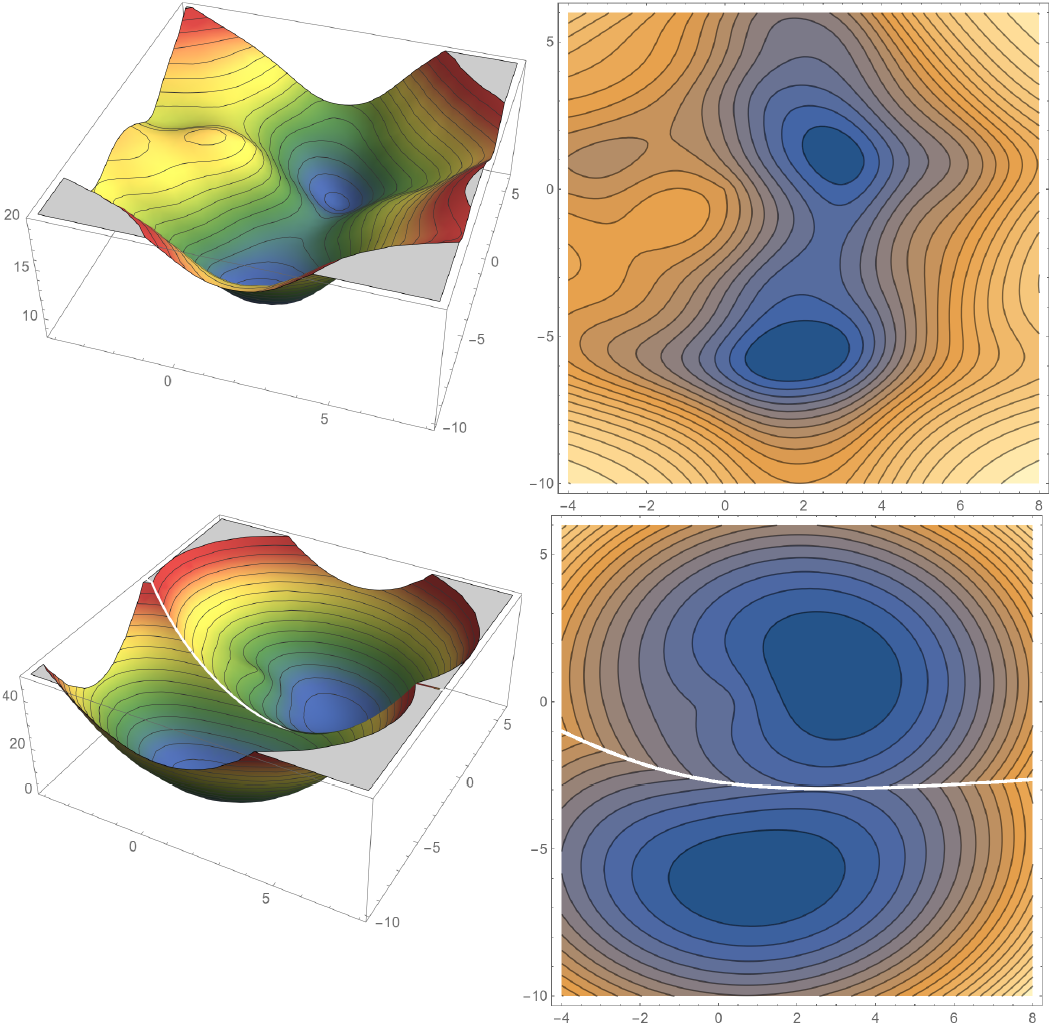
Smooth ANM minimum energy path The potential energy surface represents a hypothetical molecule with two distinct conformations corresponding to two different pairs of bond angles. The upper part shows our smooth ANM model with a genuine saddle point connecting the two energy wells, and the lower part shows the original ANM model^10^ with its characteristic singularity across the cusp (see text for details).

In addition to the path itself, the definition of the PCV also includes the mathematical formulae to measure progression along the path and the orthogonal distance from the path. There are two common types of PCVs in use in this respect, one by Branduardi and Parrinello ^11^ (BP) and one by Leines and Ensing^12,13^ (LE). We have experimented with both extensively and we primarily use the BP path, although the work presented here with the SANM path can also be applied to LE PCVs. However, the LE path requires that (1) the 3 closest path-nodes with respect to the current simulation point must be consecutive nodes, and (2) the closest node must be the middle node. In our experiences with complex biological systems, the second condition fails quite frequently, causing simulations to crash due to a discontinuity in computing the path progression. As we discuss below in Section 2 in more detail, the BP path can also exhibit discontinuities, but the barrier restraint described and implemented in this work was able to overcome those situations.

The PCV simulations in this work start with a path obtained with only C-alpha (CA) atoms. However, we found that CA-based PCV can be misleading in conformational transitions where certain side-chain perturbations are biologically relevant (e.g. the methionine flip in our STING work^14^). This shortcoming of the CA-PCV prompted us to develop a 3-stage strategy whereby we first build an approximate backbone plus C-beta (CB) chain from the CA-PCV. We then build an all-atom model on each node via a cascading minimization method outlined in Section 2.3 below. Finally, in order to refine the all-atom PCV, we run multiple simulations using this PCV, then reconstruct the PCV using selected snapshots from these simulations. This adaptive approach produces a new PCV consisting of more realistic nodes representing conformations from high-quality MD simulations. This iterative refinement procedure can be repeated to further optimize the path.

A crucial aspect of the meta-eABF procedure that we describe here is ensuring that the simulation stays close to the path (see details in the ReBaCon Section 2.4 below). In other words, the simulation should be restrained to stay within a thin tube around the path, allowing for a limited amount of sampling in orthogonal directions to the PCV. The most common restraining algorithm used in MD today is a simple quadratic penalty function such as *F*_*c*_(*x* − *x*_0_)^2^, where *x* is the variable to be restrained, *x*_0_ is its desired value, and *F*_*c*_ is a force constant. In this form the restraint penalty gets applied for all non-zero values (i.e. *x < x*_0_ or *x > x*_0_) and the restraint force is proportional to |*x* −*x*_0_|. Often we want to restrain a variable not to a particular value, but just want to make sure that the value does not exceed an upper limit and/or fall below a lower limit. For example, we want to apply an upper limit to the radius of the tube that imposes minimal bias to the simulation when close to the PCV while still limiting the search space. Historically, limits like this have been achieved by zeroing out the left hand side of the restraint parabola located at the upper limit and/or zeroing out the right hand side of the restraint parabola placed at the lower limit, resulting in the ubiquitous “flat bottom” restraint that penalizes values under the lower limit and above the upper limit whereas no penalty is applied between the limits. Mathematically, “zeroing out” means multiplying the penalty function with a unit step function centered at the upper/lower limit, respectively. While the flat-bottom potential achieves the goal of minimal bias while close to the PCV, the nature of the quadradic function has limitations, as described below.

Simple quadratic penalties as well as complex penalty functions involving higher powers, scaling, and offsets are used routinely in MD simulations. Most penalty-based restraints (e.g. polynomials) do not provide hard limits, which can allow systems to deviate from the desired region. For many situations this approach is acceptable, but not in all cases, i.e., if we want a restraint that cannot tolerate any violation. One well-known example is restraining covalent bonds involving H atoms to stay completely rigid in MD simulations to extend the time step beyond 1 fs without using multiple time step integration techniques.^15,16^ Software tool kits like Colvars^5^ and PLUMED^17–19^ allow users to define non-polynomial restraints of virtually any mathematical form. However, we are not familiar with previous publications that describe the implementation and use of barrier restraints as described in this work.

Barrier functions such as −*log*(*x*_0_ − *x*) or 1*/*(*x*_0_ − *x*) are staples of the so-called “interior-point methods” in constrained nonlinear optimization.^20^ The benefit of a barrier potential in the context of MD is that the system cannot escape because the barrier is infinite at the specified limit, a desired consequence of the asymptotic behavior. Unfortunately, interior-point methods have two significant drawbacks that makes them difficult to use in optimization problems. First, it can be nontrivial to move the constrained variable inside the allowed domain while maintaining reasonable energetics for complex biological motions. Second, and of more practical importance, is that general purpose line search methods can probe the exterior of the allowed domain where the barrier function either does not exist (logarithmic) or manifests a numerical “black hole” (reciprocal). The latter happens when the value of the escaped *x* gets infinitesimally close to the barrier wall from the outside where the barrier function goes to minus infinity. Modified line search methods do exist, but they only work efficiently with linearly constrained problems. ^21,22^ Therefore, in most cases constrained nonlinear optimization problems are solved with penalty functions utilizing a series of successively increasing force constants,^20^ but that would be impractical in MD. The basic tenet of the work presented here is to demonstrate that reciprocal barrier methods can be implemented in a way that overcomes the aforementioned issues in path-based enhanced sampling MD simulations of complex biological motions.

## 2. Methods

We ran our simulations on Nvidia RTX 2080Ti and A40 GPU-equipped, Ubuntu 20.04 workstations, using the OpenMM-PLUMED software,^17–19,23^ versions 7.7.0 and 2.8.1, respectively. The barrier function was defined in PLUMED as a custom bias. System preparation was completed with AmberTools, ^24,25^ ff14sb^26^ force field was used with proteins and gaff^27^ force field with organic ligands. All simulation job files for the work presented here have been deposited to the PLUMED-NEST repository^28–30^ and the path generator software to GitHub. ^31^ In addition, we posted and included in the Supplementary Information high-resolution trajectory movies of one of our Cullin-RING simulations^32^ (see Section 3.1) and stimulator of interferon genes (STING) simulations^33^ (see Section 3.3).

### 2.1 Well-tempered meta-eABF

For reasons that we explain below, we employ a combination of well-tempered metadynamics and extended-Lagrangian adaptive biasing force (meta-eABF) method^34–36^ for biasing the simulation along the path and estimating the potential of mean force (PMF). Consistent with the family of ABF methods, meta-eABF simulations utilize adaptive free energy biasing force to enhance sampling along one or more collective variables (CV). Meta-eABF evokes the extended Lagrangian formalism of ABF whereby an auxiliary simulation is introduced with a small number of degrees of freedom equal to the number of CVs, and each real CV is associated with its so-called fictitious counterpart in the low-dimensional auxiliary simulation. The real CV is tethered to its fictitious CV via a spring (typically with a large force constant) and the adaptive biasing force is equal to the running average of the negative of the spring force. The biasing force is only applied to the fictitious CV, which in turn “drags” the real simulation along the real CV via the spring by periodically injecting the instantaneous spring force back into the real simulation. Moreover, the main tenet of the meta-eABF method is that meta-dynamics (MtD) or well-tempered metadynamics (WTM) is employed to enhance sampling of the fictitious CV itself. The combined approach provides advantages over either MtD/WTM or eABF^37^ alone. We also demonstrate the utility of running multiple independent simulations and provide rigorous error estimates for our free energy predictions based on a boot strapping procedure.

In terms of path CV calculations described below, a path is represented by two degrees of freedom. One CV is the *S* variable representing the progress along the path and the other CV *Z* represents the orthogonal distance from the path. If we put an upper limit to *Z*, the path can be visualized as a tube with length *S* and width *Z*. The essence of modeling the transition along the path with meta-eABF is biasing either *S* (1D bias) or both *S* and *Z* (2D bias) to produce a simulation that follows the path back and forth along *S* while staying inside the tube, or actively exploring the the *Z* dimension as well in case of a 2D simulation.

### 2.2 C-alpha path generation

We employ a 3-stage minimum-energy path generation strategy that extends Roux’s anisotropic network model (ANM).^10^ The stage 1 C-alpha path-CV (CA-PCV) works for many biological systems, but we discovered that CA-PCV can be misleading in conformational transitions where, e.g., certain side chain conformational changes are biologically relevant but have barriers such that they are not sampled sufficiently without enhanced sampling. Simulations using a CA-PCV can readily transform the CA trace between two protein conformations, but the critical side-chain transformations may not follow, which can prevent the simulations from reaching the true biological endpoints. For systems like this, it is helpful to include certain side chain atoms in the path to provide more biasing leverage. Our strategy continues with stage 2 where we first build a backbone plus C-beta (BB+CB) path based on simple geometric considerations, followed by stage 3 where we build an all-atom (AA) path employing a cascading implicit-solvent minimization method (see below).

The ANM model consists of two separate

Gaussian network models centered at conformation A and conformation B, respectively. A harmonic potential with *F*_*c*_ = 1 is applied to all pairs of CA atoms at a distance *<*= 8 Å, and *F*_*c*_ = 0 for all other pairs. The pairwise potentials are summed up separately, for conformation A (ANM-A) and conformation B (NMA-B), respectively, and the potential energy function of the combined ANM model is MIN(ANM-A, ANM-B). ^10^ This can be visualized in 2D as shown in the lower part of Figure 1 with the white gap representing the non-differentiable singularity where ANM-A = ANM-B. Roux’s algorithm locates the approximate minimum along the singularity and from there initiates two separate minimization paths toward minimum A and minimum B, respectively. The minimum potential energy path (MPEP) is constructed by concatenating the A and B paths.

While the above ANM model has utility, it suffers from a discontinuity at the cusp, where the two Gaussians intersect. To overcome this, we propose a smooth ANM (SANM) model that has no singularity and is based on the combination of two network-like models such that the potential energy function is a “sum” instead of “min”—SUM(SANM-A, SANM-B). The upper part of Figure 1 shows an example of SANM model, which contains no singularity—indeed, the SANM produces a genuine saddle point connecting the two minima. Note that the corresponding ANM and SANM minima do not coincide exactly, but the difference is small in our experience, typically less than 0.5 Å C-alpha root-mean-square distance (CA-RMSD), which is well within thermal fluctuations at room temperature. The functional form of SANM can be virtually anything that approximates the quadratic at the minimum, but is flattened out further away from the minimum. We use the following function in Equation (1), where *d*_*ij*_ is the pairwise CA-CA distance in any conformation while 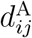 and 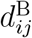are the same in the reference conformations A and B, respectively. Only a subset of atom pairs *i* and *j* are included in the sums for which the arithmetic mean of 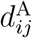and 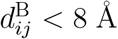. We refer to the left sum as SANM-A and the right sum as SANM-B. In the vicinity of conformation A SANM-A is near quadratic while SANM-B is approximately flat (equal to the number of *i, j* pairs included in the sum), and vice versa in the vicinity of conformation B. Figure 1 (top) gives a good visual representation of Equation (1) in two dimensions.

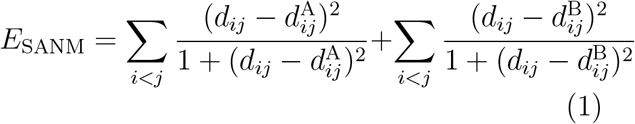

In principle we could generate a CA-path on the SANM potential energy surface (PES) by locating one or more saddle points about halfway between the two minima. Although this approach seems viable, finding saddle points on a multidimensional PES is very inefficient compared to minimization. Given our objective to have a method that can be applied in real-world drug discovery projects fast enough to make an impact on decisions, we favored a practical approach that involves only minimization. We start by minimizing the SANM energy of one of the endpoint structures, which is usually an X-ray structure, and arrive at one of the minima on the SANM-PES, say A. This is the first short segment of the CA-path. As we stated above, the minima on the SANM surface are within 0.5 Å RMSD from their corresponding physical structure in our experience. A similar terminal segment is then added to the path by minimizing the other endpoint structure and finding the other minimum B. The significant middle segment of the CA-path is generated by restrained minimization. Starting from minimum A *E*_SANM_ is minimized subject to position restraints (POSRE) representing minimum B. The key for this algorithm to work is to use a weak restraining force constant. We found that 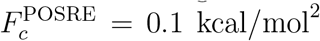 was particularly efficient in gently pulling the minimization from one minimum to the other, passing a low energy barrier on the *E*_SANM_ PES. This barrier is generally not a true saddle point, but works well for all practical purposes. As a consequence of the above procedure, the algorithm generates two different CA-paths for a given pair of terminal segments, depending on whether we start in minimum A or minimum B. Unless the exact nature of the transition state is important, either path can be used to estimate the PMF, but using both independently can further enhance sampling and provide a better estimate.

The SANM model presented here extends the ANM model in another aspect as well. Specifically, we noticed that the standard ANM model without additional restraints exhibits an undesirable artifact that manifests in strongly curved, tight transition paths, as is often the case in highly coupled condensed phase systems such as proteins in water. The ANM model works well in the vicinity of the reference conformation but long amplitude excursions (as is often needed to simulate biologically relevant motions) can result in nonphysical distortions in the geometry. For example, using solely the ANM model can produce a CA-path where consecutive node structures in a flexible loop fold on top of each other and some CA atoms nearly overlap. Therefore, we included three additional restraint terms in the SANM model. CA-CA virtual bonds are kept at approximately 3.8 Å distance, (2) CA-CA-CA angles are restrained to 75−145°, and (3) non-adjacent CA-CA pairs are also kept apart at least 3.8 Å. With these restraints even the tightest turns in the CA-path maintained physical geometries.

### 2.3 Backbone and all-atom path

For systems where side-chain packing is an integral part of a conformational transition, the CA-path itself is often inadequate because the side chains do not properly reorient during the transition. It is possible to build an all-atom (AA) path but it would be inefficient from a computational perspective to use all the side-chain atoms—usually a small subset will suffice. The bottleneck in path calculations is twofold, (1) passing the atomic coordinates between the MD engine and the advanced sampling module at every time step, plus (2) computing RMSD to multiple path nodes. Theoretically, these operations scale as *O*(*N*_at_) and *O*(*N*_at_log(*N*_at_)), respectively, at best and 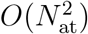 at worst, with the number of atoms *N*_at_ included in the calculation. Moreover, state-of-the-art MD engines are all GPU based, but the path-CV/meta-eABF modules^5,17–19^ run on multiple CPUs and, therefore, even a modest reduction in *N*_at_ can result in significant overall simulation speed-up, so it is highly beneficial to minimize *N*_at_.

The first step towards the AA-path is the construction of the backbone and CB atoms on top of the CA trace (BB+CB-path). The BB+CB-path need not be highly accurate because it is not used here directly for free energy estimations—we just utilized the lever effect of the CB atoms to achieve the correct side-chain transitions when it was necessary. We employed a simple algorithm based on geometric considerations. ^31^ Assuming that all-atom models of both path endpoint conformations are available, constructing the AA-path continues as follows. The number of nodes *N* is the same in the CA, BB+CB, and AA paths, respectively. (1) First, we replace the CA and CB atom coordinates of conformation A, which is AA-path_1_ with those of the adjacent node in the BB+CB-path BB+CB-path_2_, and minimize the energy using an implicit solvent model while restraining the CA and CB coordinates to their new values. The resulting minimum energy structure has approximately the same CA and CB coordinates (depending on the POSRE force constant, namely how much relaxation we allow) as in the corresponding BB+CB-path_2_ node and the side chains will be slightly different from their original configuration in AA-path_1_. This structure is now AA-path_2_ and serves as the basis for generating AA-path_3_. (2) Then, we replace the CA and CB coordinates in AA-path_2_ with those in BB+CB-path_3_ and apply the same POSRE minimization protocol to construct AA-path_3_. (3) Next, we continue the cascading procedure until we reach AA-path_*N/*2_ and then start over from the other end. (4) Finally, starting with conformation B, i.e., AA-path_*N*_ we continue the cascade backwards until we reach AA-path_*N/*2+1_ and at that point the full AA-path is completed. After pruning the AA-path to include only relevant side-chain atoms, this path-CV can be readily used for simulating complex protein conformational transitions.

The final contribution that we present in this work is an improvement in the structure of the nodes used along the path. The nodes of the path described above consist of minimized structures in implicit solvent, but it would be more accurate if the nodes represented snapshots (saved trajectory frames) from a high-quality simulation with explicit solvent. First, we run multiple independent meta-eABF (or any other kind of enhanced sampling) simulations using the pruned AA-path for a few hundred ns, saving trajectory frames quite frequently (approximately every 50-100 ps). Then, from the combined pool of trajectory frames we select a subset such that it best resembles the original path, but now the nodes are replaced by frames from an all-atom explicit solvent molecular dynamics (MD) simulation. We used the following simple algorithm for reconstructing the AA-path. (1) Each original node is assigned a set of frames from the combined trajectory for which the RMSD from that node is less than a user-defined cut-off value, typically *<* 0.5 Å. Note that there will be significant overlap among the S_1_-S_*N*_ sets, but that is not a problem. (2) The first node of the new trajectory-frame based path can be randomly selected from any of the S_1_-S_*N*_ sets. Here we select a frame from S_1_ to initiate the new path with AA-traj-path_1_. (3) AA-traj-path_2_ is selected from S_2_ but the selection is no longer random, instead the frame in S_2_, which is the closest in RMSD to AA-traj-path_1_, is the one selected. (4) AA-traj-path_3_ and so on are added to the new path in a similar fashion, AA-traj-path_*i*_ is selected from S_*i*_ adding the shortest segment to the path in terms of RMSD measured from AA-traj-path_*i−*1_. Adding the shortest segment instead of random selection helps maintain fairly evenly distributed nodes along the new path, which is an important requirement in both BP^11^ and LE^12,13^ paths, especially the latter. We note, however, that enforcing strictly evenly spaced nodes can introduce artifacts in terms of non-physical geometry and this is another reason we prefer the BP path, which is tolerant to slight variations in node spacing, while the LE path is very strict about keeping even spacing. Also note that each node in AA-traj-path must be unique (already selected trajectory frames are removed from the pool during path reconstruction) to assure monotonic RMSD change along the path with respect to the endpoint conformations A and B. (5) This procedure can be iterated to further improve the path by running additional simulations with the new path and collecting more trajectory frames.

### 2.4 Reciprocal barrier restraint (ReBaCon)

In order to introduce the new barrier restraint we first provide a closer look at the traditional approach. As an example, we show an upper wall restraint located at 1 (i.e. we do not want variable *x* to go above 1). This upper wall is represented as a black vertical line in Figure 2. On the top right we show four different penalty functions using powers 2 to 5 (blue, orange, red, and green, respectively), and for simplicity we choose *F*_*c*_ = 1. Θ is the unit step function. The traditional quadratic penalty is represented by the blue curve. There are two features to note: (1) the penalty function only couples with the system for *x >* 1, and (2) these kind of penalty functions have very slow acceleration and therefore grow very slowly. In fact, the faster, higher powered functions start making a significant difference only for *x >* 2. Penalty functions such as this are only good for weak restraints in MD simulations. The negative derivative of the penalty function/potential is the restraining force and its magnitude is proportional to the restraint violation. Figure 2 (bottom right) shows the forces associated with the penalty potentials. The forces are negative, pulling *x* back toward 1.

**Figure 2:**
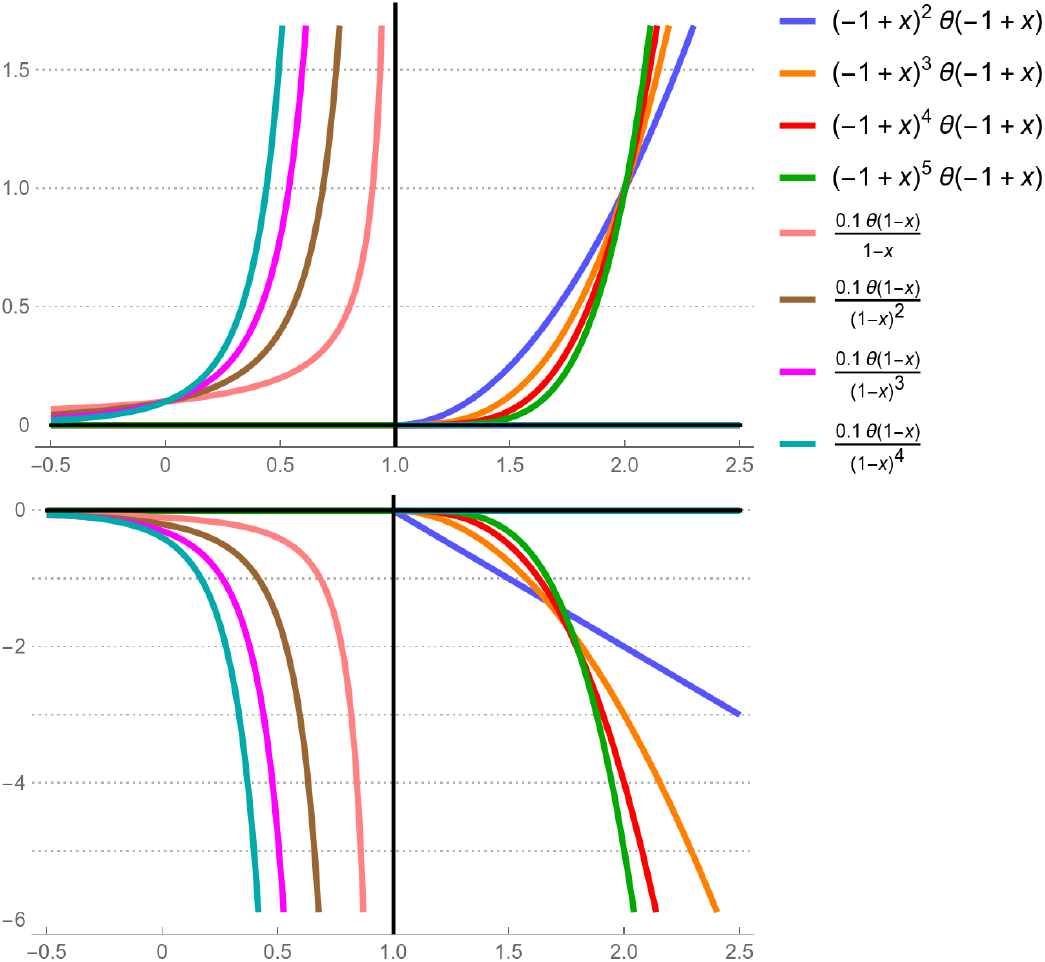
Traditional polynomial penalty versus reciprocal barrier constraint. The upper part shows the restraint potential of a series of barrier restraints (left) and penalties (right), and the lower part shows the corresponding restraint forces. The upper wall is represented by the vertical black line located at 1.0, see text for details.

From a practical perspective, it would be preferable for the restraint potential/force to engage inside the wall and not outside. In that spirit, barrier functions are designed to never let the restrained variable escape from its allowed domain by applying an infinite wall. Figure 2 (top left) shows a series of such barrier functions that are actually related to the penalty functions, but instead of using positive powers, they use negative powers. In general, we apply a simple and robust barrier function of the form 1*/*(*x*_0_ − *x*), which is used in all of the path simulations presented in this work, although for visual purposes the barrier functions are scaled by 0.1 in Figure 2.

Barrier functions are never zero in the allowed domain, but their values are very small up until they get close to the wall, where they grow exponentially fast and go to infinity in the limit. Higher negative powers present a steeper wall and allow *x* to get closer to the wall. Similar effect can be achieved with the lower powers by multiplying them with a small constant *k <<* 1. However, in MD simulations there is a limit to this, in our hands *k <* 0.008 kcal/mol^2^ resulted in occasional breaks through the barrier wall due to the finite MD time steps. Figure 2 (bottom left) shows the barrier forces. Again, the forces are negative, pulling *x* away from the wall, keeping it inside. Also note how much larger the magnitudes of the barrier forces are in the vicinity of the wall with respect to the penalty forces on the bottom right. In the Results we provide specific reasons why barrier restraints are superior to penalties for specific examples.

## 3 Results and Discussion

In this work, we describe a novel method for enhanced sampling MD that combines a smooth ANM (SANM) path collective variable (PCV) with a barrier restraint to efficiently simulate complex conformational events in biological systems. Specifically, we study three diverse systems with different structural compositions and conformational mechanisms. In the first example, we simulate the motions in the Cullin-RING ligase (CRL), which is responsible for ubiquitination for many E3 ligases involved in targeted protein degradation (TPD), with von Hippel-Lindau (VHL) as the E3 ligase and SMARCA2 as the protein of interest. Next, we study the active/inactive transition involved in oncogeneic JAK2, where the V617F mutation in a non-ATP pocket of the pseudokinase domain results in hyperactivity of the kinase. Finally, we explore the conformational landscape of a particular loop in STING (Stimulator of Interferon Genes) that we found to be important during our discovery and development of SNX281, a novel small molecule STING agonist in the clinic. The breadth and complexity of these systems offers a good case study for conformational free energy simulations and the results demonstrate the robustness of the novel approach presented in this work.

### 3.1 Cullin-RING Ligase with a VHL-degrader-SMARCA2 Ternary Complex

Heterobifunctional degraders, which consist of two separate protein binding moieties (the warhead and the E3-ligand) joined by a linker, are a class of molecules that “induce proximity” between a target protein of interest (POI) and an E3 ubiquitin ligase. This induced proximity can lead to ubiquitination of the POI and its subsequent proteosomal degradation. Targeted protein degradation (TPD) presents a novel approach to drug protein targets, since a single degrader molecule can induce catalytic degradation of the POI and potentially offer an avenue to eliminate targets traditionally labeled as undruggable by classical therapeutic strategies. In the work presented here, we assemble the entire eight-protein Cullin-RING ligase (CRL) supramolecular complex to explore structural and dynamic factors associated with ubiquitination. Specifically, we probe the conformational landscape associated with different solvent-exposed POI lysine residues coming into proximity with the E2-loaded ubiquitin of the CRL, specifically focusing on the probability of POI lysine residue density within this “ubiquitination zone”. We recently published a broad study on atomic-resolution predictions of degrader-mediated ternary complex structures,^38^ which included a preliminary study of the dynamic nature of the full CRL complex via meta-eABF. However, in that work we encountered a problem with traditional penalty restraints, which prompted research into a better solution and resulted in the barrier restraint described in this work.

Figure 3 shows the individual nodes representing discrete conformations along the path. The fully open and closed conformations of the CRL complex were constructed based on partial experimental structures (PDB IDs 1LQB, 5N4W, and 6TTU) as we described in detail elsewhere. ^38^ The gradually morphing conformations are shown as smooth CA-traces colored by a red-to-white-to-blue palette, red representing the closed conformation and blue representing the open conformation. For reference, we show each of the biological units comprising the supramolecular complex as colored patches on the composite surface representation that includes both open and closed conformations. One can visualize the path CV as “morphing” between the two endpoints, but the minimum potential energy path (MPEP) is significantly more realistic than a direct geometric morph.^10^

**Figure 3:**
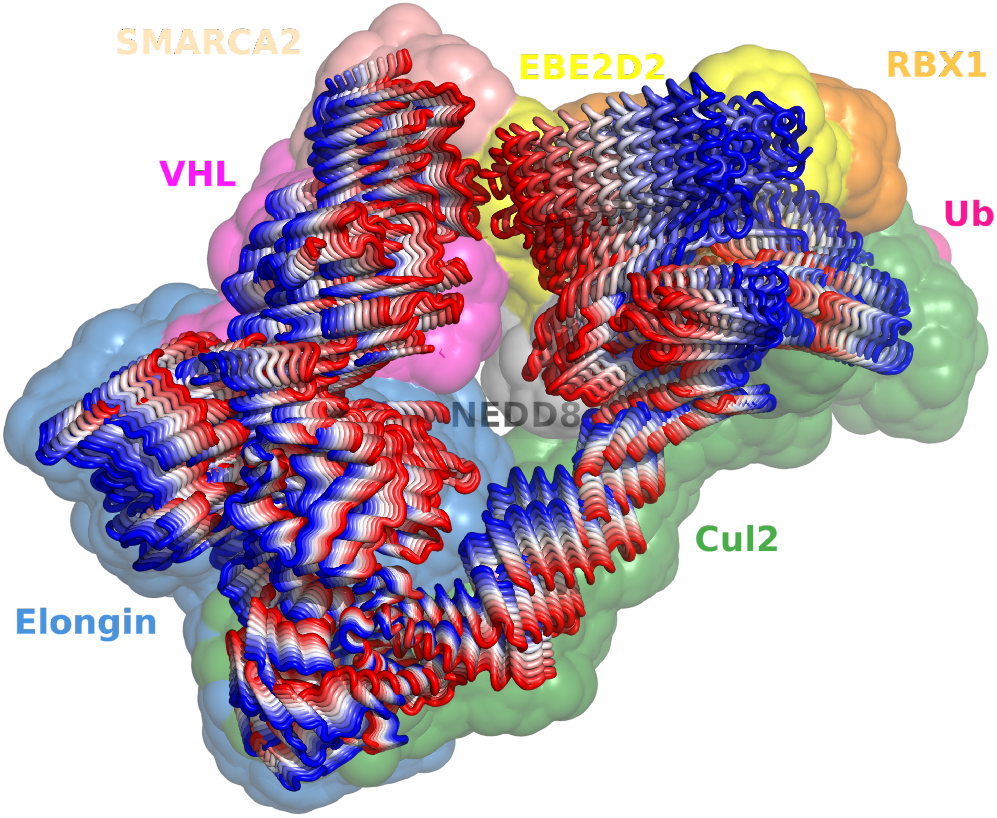
Frame stack representing the path collective variable (PCV) for the open-to-closed transition in the full CRL-VHL-degrader-SMARCA2 complex. The individual conformations along the path are shown as CA-traces colored by a red-to-white-to-blue palette, red representing the closed conformation and blue representing the open conformation. The biological units that comprise the CRL complex are shown as colored patches behind the stacked frames. A movie of one of our open/closed transition simulations is provided in the Supporting Information.

Computation of the PCV in a system of this size undergoing a non-linear conformational motion is quite complex^11^ even when it is represented by only two degrees of freedom. One CV is the *S* variable, representing the progress along the path (e.g., if we have 50 nodes then *S* goes from 1 to 50). The other CV (*Z*) represents the orthogonal distance from the path. When we apply an upper limit to *Z*, the path can be visualized as a tube with length *S* and width *Z*. The essence of modeling the open and closed transition, then, is running a meta-eABF simulation such that the *S* collective variable follows the path back and forth and the *Z* variable keeps the system within a specified distance of the path, allowing sampling with minimal bias within the tube. This scenario is where the new kind of restraint presented in this work shows its strength.

It is instructive to note that the maximum radius of the path tube (*Z*_max_) should be small enough to constrain the conformational space while being large enough to allow unbiased sampling orthogonal to the path. In our experience with protein systems, this varies between as thin a tube as *Z*_max_ = 0.2 Å radius to about 3.0 Å, depending on the system and the path. There are two independent reasons why this limit exists. (1) The BP path algorithm is exact only for thin tubes^11^—beyond that the computation of *S* and *Z* becomes indeterminate and causes jumps in the *S* and *Z* variables from one MD time step to the next. (2) In case of highly curved paths associated with complex conformational transition patterns, a simulation that strays too far from the center of the path can jump from one part of the *S* path to another in a non-continuous way. To visualize this, consider a PCV shaped as the letter Ω. As the orthogonal distance *Z* increases (i.e. the thickness of the line), the bottom-left and bottom-right of the Ω will eventually overlap. As such, a simulation with a large *Z* can inadvertently jump from the left base to the right base, completely bypassing the entire curved part of the path (the original intention of the PCV). With a smaller *Z* the simulation can stay on the path throughout its full length.

The problem with jumps in CV values is that they cause discontinuities in the spring force coupling the real CV and the fictitious CV, which will ultimately result in the simulation crashing. To avoid these discontinuities, we found (by trial and error) that the tube radius should be 3.0 Å for the CRL-VHL-degrader-SMARCA2 complex. While we could use a penalty restraint to achieve this, we show that using a reciprocal barrier restraint has benefits. For example, the standard harmonic penalty applied at 3.0 Å distance would require an offset (explained in section 2.4) with the penalty starting at *Z* = 0. Using this offset we would need a fairly large force constant *F*_*c*_ ≈ 1, 000 kcal/mol/Å^2^ to keep the simulation inside the 3.0 Å tube, which is large but still practical. However, if we set the maximum radius to 0.5 Å we need a *F*_*c*_ ≈ 100, 000 kcal/mol/Å^2^, which can be prohibitive in MD simulations. Regardless of whether or not it is feasible to use a particular large force constant, the problem with a penalty is that the concrete value of the required *F*_*c*_ depends on the nature of the free energy surface and the tube radius, which can only be determined empirically based on numerous test simulations on a particular system. Conversely, the exact same reciprocal barrier function 1*/*(*R*_max_ − *Z*) can be used for any tube radius. It takes exactly the same amount of barrier restraint force to enforce a 3.0 Å tube radius as it takes to enforce a 0.5 Å tube radius.

In order to demonstrate the adverse effects of using inadequate harmonic restraint and show the benefits of the reciprocal restraint, we ran meta-eABF simulations using the harmonic restraint *F*_*c*_*Z*^2^ set at *Z* = 0 with *F*_*c*_ = 250 kcal/mol/Å^2^ and the barrier restraint *k/*(3 − *Z*) where *k* = 1 kcal/mol*Å and the outer surface of the tube is set to be at *Z* = 3Å distance from the path. In Figure 4 we compared the restraint potentials and forces for barrier vs. penalty at two levels of magnification, wide range (top) and in the close vicinity of the tube surface (bottom). The horizontal axis represents the tube radius *Z*. The positive domain of the vertical axis shows the restraint potential in kcal/mol and the negative domain represents the restraint force in kcal/mol/Å. The restraint force is the manifestation of the wall and as Figure 4 shows, the barrier force approximates a hard wall whereas the penalty force represents a soft wall. The orange curve is the harmonic penalty potential with *F*_*c*_ = 250 kcal/mol/Å^2^ with its value equal to 2,250 kcal/mol at *Z* = 3 Å, and the green curve is the corresponding linear penalty force. The harmonic potential grows slowly, making it ineffective at keeping the system within the desired region. This is vividly demonstrated at high magnification shown in the bottom part of Figure 4.

**Figure 4:**
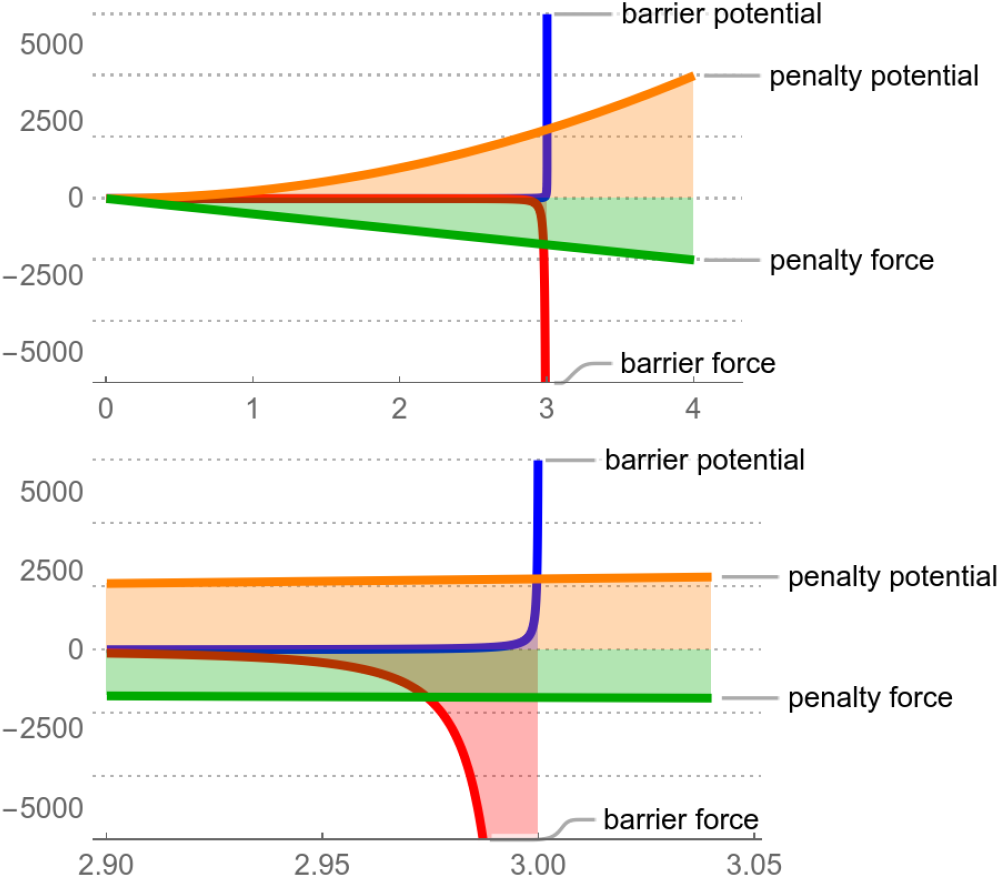
Path tube restraint: barrier vs. penalty Comparison of a traditional harmonic penalty with the reciprocal barrier restraint presented in this work. Two levels of magnification are shown, wide range (top) and in the close vicinity of the tube surface (bottom). The horizontal axis represents the distance *Z* from the center of the tube. The positive domain of the vertical axis shows the restraint potential in kcal/mol, and the negative domain represents the restraint force in kcal/mol/Å. The orange curve is the harmonic penalty potential and the green curve is the corresponding force. The penalty starts at *Z* = 0 and therefore has a substantial value close to the wall, but as the insert in Figure 6 (purple curve) shows, the simulation still proceeds outside the wall. In stark contrast, the barrier restraint (blue curve potential, red curve force) is negligible all the way up to the immediate vicinity of the wall where it provides an infinite barrier preventing the simulation from ever leaving the tube.

Figure 5 shows the evolution of the *S* variable in a slice of the meta-eABF simulation of the CRL-VHL-degrader-SMARCA2 system from 30-60 ns. The vertical axis shows the index of the path CV nodes as units, which roughly corresponds to Angstroms in this case. Values close to 1 represent the closed conformation and close to 40 representing the open conformation. First, consider the purple plot, which corresponds to using a traditional harmonic restraint on *Z*. It exhibits several spikes where *S* jumps several units, showing evidence of discontinuity. The simulation survives for a while but eventually crashes at about 55 ns. The insert in Figure 5 displays a magnification of the middle part to show the spikes more vividly between 43 ns and 47 ns.

**Figure 5:**
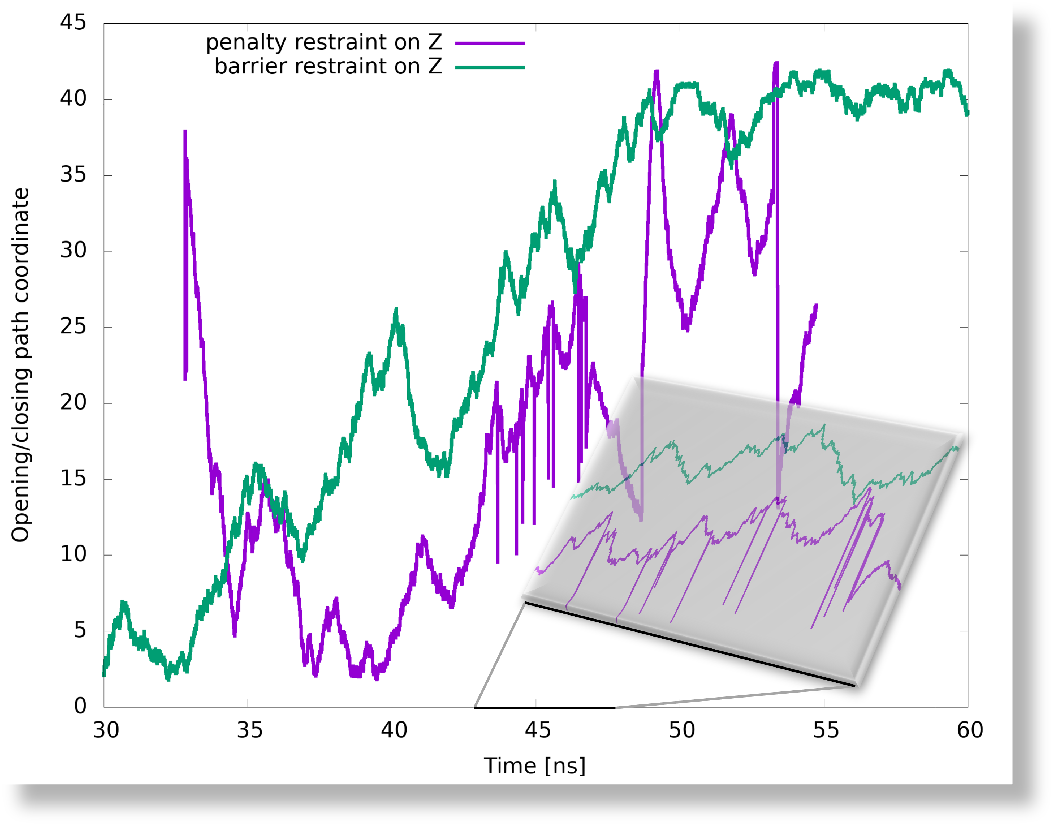
Time evolution of the *S* path variable using a harmonic penalty restraint on the *Z* variable (purple) shows significant jumps along the path shown by the spikes along the vertical *S* coordinate, and crashes at about 55 ns. However, using the reciprocal barrier restraint (green), the simulation is smooth and runs indefinitely, see text for details. The insert shows a magnified view between 43 ns and 47 ns.

Figure 6 shows the same issue in the *Z* dimension. The purple spikes are problematic—as mentioned above, the *S* and *Z* spikes originate from a mathematical formula^11^ that breaks down at about *Z* = 3.0 Å with this system. While the spikes are clear visual cues, the critical data in Figure 6 are the values of *Z* where there are no spikes, between 34 and 43 ns, as shown in the insert where the “normal” purple *Z* values hover between 3-4 Å, which is beyond the desired tube radius of 3 Å.

**Figure 6:**
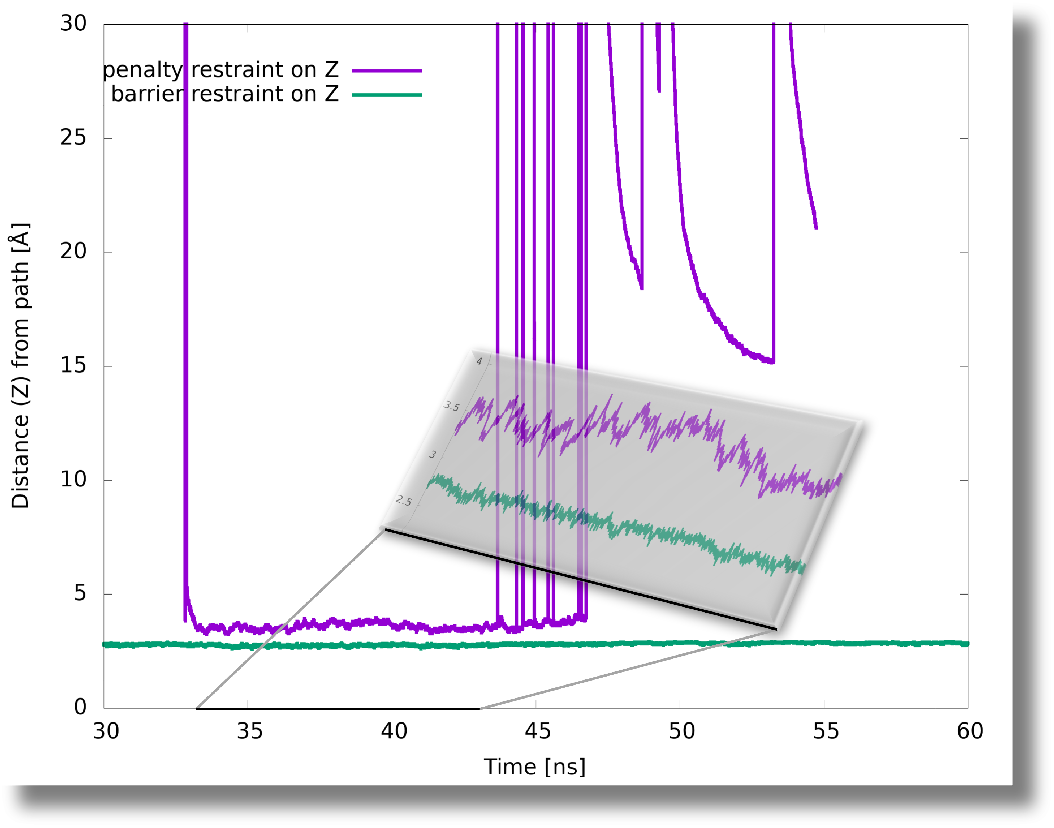
Time evolution of the *Z* path variable. The jumps in the *S* variable shown in Figure 5 are coupled with enormous jumps in the *Z* variable (purple spikes) whereas the reciprocal barrier restraint keeps the *Z* value safely below 3Å (green curve; see text for details). The insert shows a magnified view between 34 ns and 43 ns.

Now, consider the green plots representing the exact same simulation except for using the barrier restraint in place of the harmonic penalty. Here, both *S* and *Z* are smooth (the plots show data points at 1,000 MD time step intervals), *Z* stays below 3.0 Å, and the simulation runs without incident for the specified time of 100 ns. Since the penalty starts at *Z* = 0, the harmonic penalty has a substantial value even when relatively close to the wall, but as the insert in Figure 6 (purple curve) shows, the simulation still proceeds outside the wall (up to 4.0 Å from the path. In contrast, the barrier restraint (blue curve potential, red curve force in Figure 4) has negligible value all the way up to the immediate vicinity of the wall where it provides an infinite barrier preventing the simulation from ever leaving the tube. In fact, with *k* = 1 kcal/mol*Å the barrier restraint kicks in much earlier and as shown in the insert in Figure 6 (green curve), the simulation stays within ∼2.8Å of the path. From the above example, the benefits of the SANM path coupled with the novel barrier restraint allow for efficient and robust sampling along the path, which was not possible with the traditional ANM path and harmonic restraint.

### 3.2 JAK2-V617F pseudokinase domain

The oncogenic JAK2 V617F mutation lies in the pseudokinase domain of JAK2, which is distal from the pseudokinase ATP binding site and the kinase catalytic site.^39^ This suggests that the impact of the V617F mutation might involve a conformational mechanism. Crystal structures of the V617F domain in complex with two compounds that binding in the ATP site of the pseudokinase reveal a conformation that is characteristic of the wild-type domain, rather than that previously observed for the apo V617F mutant. These structures suggest that certain ligands in the ATP site of the pseudokinase can modulate the V617F mutant and restore a wild-type conformation, thereby providing a rationale for the allosteric mechanism.

Here, we aim to rationalize the McNally^39^ finding that Compound 2 in complex with the V617F domain (PDB code 6G3C) favors the wild-type (WT) domain conformation (PDB code 4fVQ) rather than the mutated oncogenic form (PDB code 4FVR). As shown in Figure 7 the two structures are virtually identical except for the small but crucial differences around the site of mutation. The conformational differences include a displacement of the helix and change in loop conformation connecting the helix to the beta-strand. Figure 8 shows a magnified view of the region around the V617F mutation.

**Figure 7:**
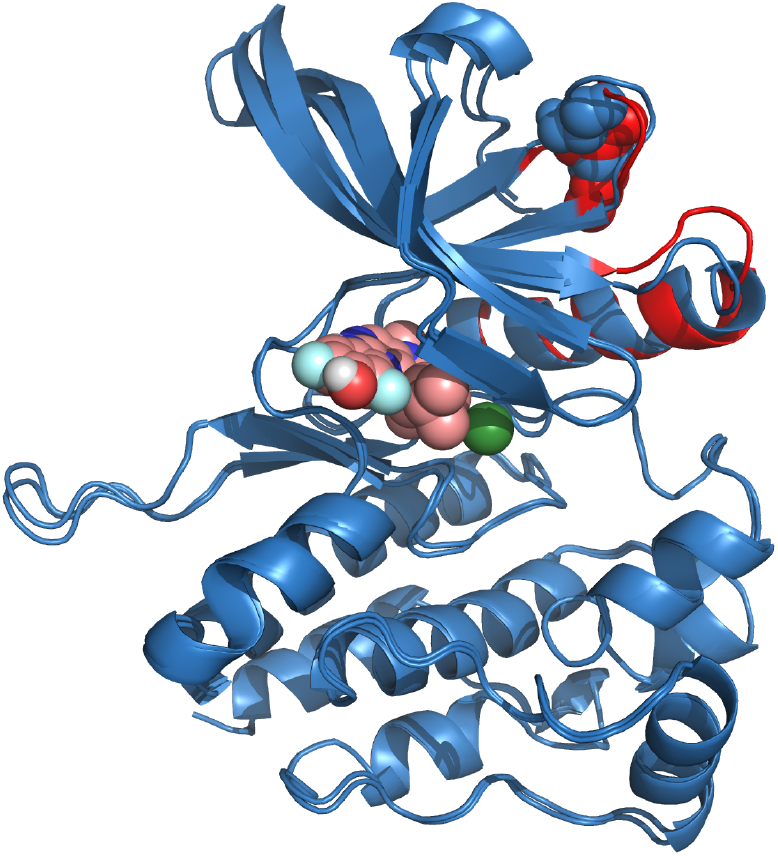
Superposed X-ray structures of the wild-type and the V617F mutant form of the JAK2 pseudokinase domain in complex with Compound 2^39^ The two structures are virtually identical except for the small but crucial differences around the V617F mutation. The wild-type (WT) conformation is red and V617F is blue. The conformational differences are highlighted in red, which include a displaced helix and loop movement. Residue F617 is shown in CPK. Compound 2 in the ATP binding site is also shown in CPK. The insert in Figure 8 shows a more detailed view of the allosteric site.

**Figure 8:**
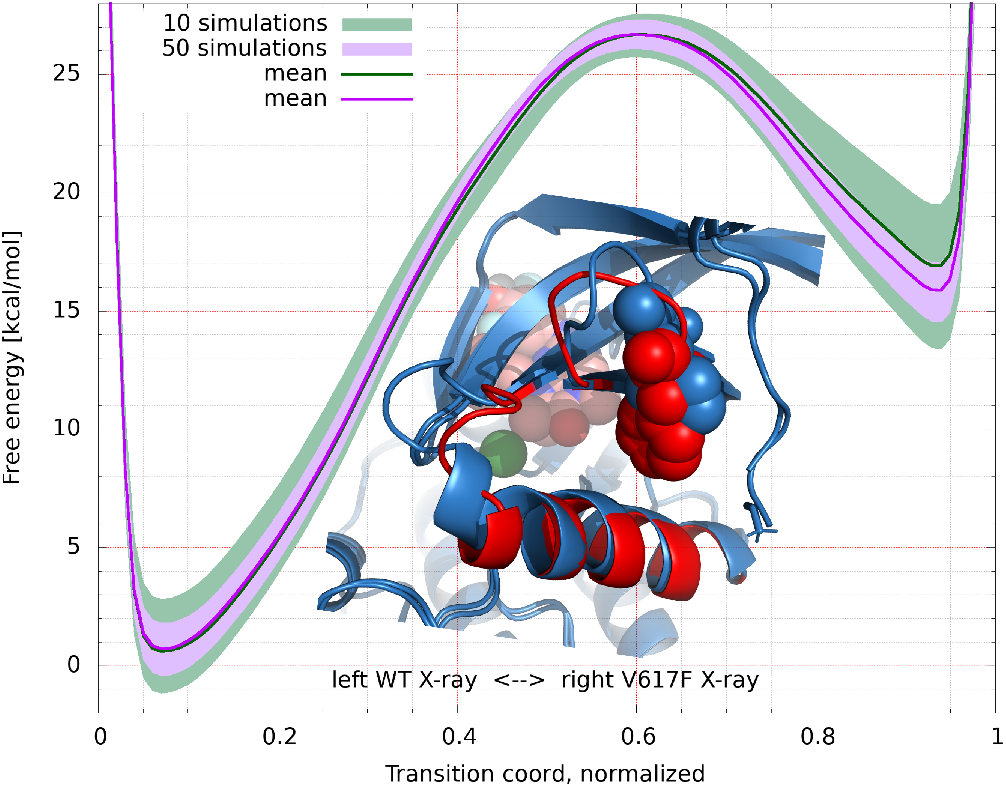
Average PMF of 1,000 boot strapped samples at 99% confidence interval. The horizontal axis represents the transition coordinate of the conformational transformation between WT (left) and V617F (right) forms of the JAK2 pseudokinase domain. The vertical axis shows the free energy in kcal/mol. The PMF plot includes the average and the associated 99% confidence interval computed from 10 (green) and all 50 (purple) independent meta-eABF simulations. The insert shows a detailed view of the conformational transition. WT conformation is red and V617F mutant is blue.

Since the conformational differences between WT and V617F away from the site of mutation are relatively small, the CA-path was chosen to only include residues near the site of mutation. We applied two-dimensional meta-eABF bias using both path parameters *S* and *Z* with a tube radius of 0.5 Å. We found that 60 ns of meta-eABF simulation produced multiple full sweeps of the path. To provide a statistical error estimate for the prediction of the conformational free energy barrier between WT and V617F, we ran 50 independent simulations starting with the same structure but assigning different random velocities from a Maxwell distribution on a per atom basis.

Our aim was to compute the free energy surface along the transition path between the WT and V617F conformations, and provide rigorous statistical error estimate. In meta-eABF the PMF is computed by numerically integrating the biasing force. In a one-dimensional simulation where only *S* is biased, the biasing force is binned along the path. Each bin contains two pieces of data, one is a counter that is incremented every time the simulation point has an *S* coordinate inside that bin, and the other is the running average of the negative spring force 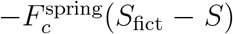 where the current value of *S* falls within the bin. This näive estimate is biased, though, and only works well for fully converged simulations. In most cases, as here too, we used the corrected z-score averaged restraint (CZAR) estimate. ^35^ In a two-dimensional meta-eABF simulation both *S* and *Z* are binned in a two-dimensional lattice, in which case the numerical integral provides a two-dimensional PMF. In this study we were only interested in the PMF along the path, so we marginalized the 2D PMF to get the 1D PMF along *S*.

The 50 independent simulations afforded 50 different PMFs, which were combined into a single PMF using boot strapping to compute a confidence interval along the entire PMF. To do this we first normalized the arbitrary integration constant *C*_*i*_ associated with every PMF_*i*_. Computing the *C*_*i*_ constants requires a boundary condition—we used Steiner’s theorem^40^ originally stated in classical mechanics (“Trägheits-Momente) but also furnishing a least square estimator in a conditional setting. In our setting, we seek a set of vertical shifts such that that overall vertical variance along the shifted PMF curves is minimized.^41^ We computed the 99% confidence interval around the mean of the optimally shifted PMF curves using 1,000 boot strapped samples of 10, 20, 30, 40, and all 50 PMFs. On Figure 8 we plotted the average PMF from 10 and from 50 simulations. The PMF indicates a fairly converged simulation with not much difference in the mean, although running 50 independent simulations significantly narrowed the confidence interval. The meta-eABF simulations are consistent with the experimental observation^39^ that Compound 2 in complex with the V617F domain shifts the conformational landscape toward the the WT conformation and away from the oncogenic form. Figure 8 also shows the highly localized conformational change between WT (left, red) and V617F (right, blue), indicating that the binding of Compound 2 in the ATP site has a significant effect in the allosteric site.

### 3.3 Stimulator of Interferon Genes (STING)

STING (stimulator of interferon genes) is a homodimer protein involved in regulation of the innate immune system and plays a role in antitumor immunity by inducing the production of cytokines. Activation of STING results from the binding of endogenous cyclic dinucleotides (CDNs) that induce a conformational change to initiate signaling. We previously published the design of a systemically available small molecule STING agonist (SNX281) that is currently in clinical trials for a variety of tumors. ^14^ Here, we provide details of conformational free energy simulations, which provide insights into the structural and dynamic basis of a specific transition observed in our drug discovery efforts.

We simulated the Met-loop conformational change to gain insight into the complex multi-step mechanism and better understand the biology related to ligand binding. The STING activation process involves multiple conformations, and in particular we were interested in the specific transition shown on Figure 9 where the large loop movements are coupled with the 180° flip of M267 from the “in” to the “out” orientation in the so-called Met-loop (residues 262-272) transition that we first noticed in our in-house X-ray structures^14^. This computational study presented challenges with a traditional ANM path and harmonic restraints, spurring us to employ the SANM path with a barrier potential. We followed our 3-stage strategy presented in sections 2.2 and 2.3. The CA-path was not suitable to model the Met-flip, so we generated the AA-path and pruned it to include the methionine sulfur atom of M267 in both units of the STING dimer. This small but significant change in the path afforded the first simulations where we observed the Met-flip (see trajectory in the Supplementary Information). For this simulation we used a thin path-tube with *Z* = 0.2 Å radius to avoid the simulations crashing for reasons described in section 3.1. We could not have achieved this using traditional penalty restraints, but as we pointed out in 3.1, when using the reciprocal barrier restraint presented here, it takes the same amount of restraint force independent of the tube radius. However, with very thin tubes the biasing force in meta-eABF tends to be very large because there is no room to relax in the orthogonal *Z* dimension and, therefore, the free energy surface tends to be inaccurate.

**Figure 9:**
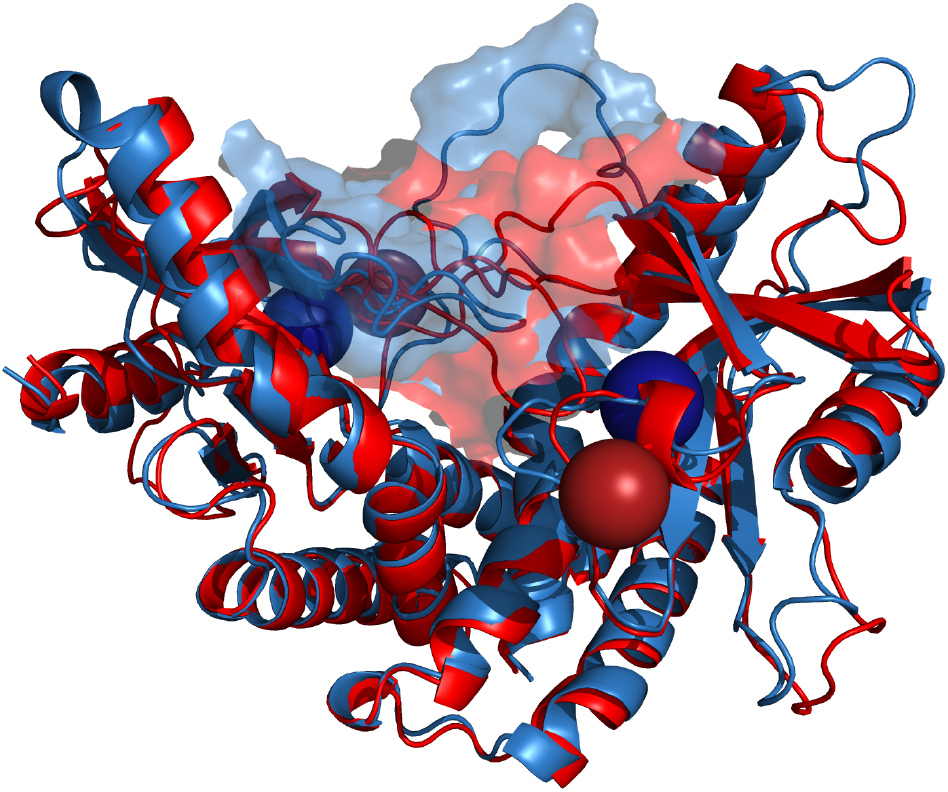
Superposition of two conformations of the STING dimer, highlighting a large change in a methionine-containing (Met) loop. The Met-in and Met-out conformations have numerous distinct differences. The Met-in conformation is shown in blue and the Met-out conformation in red. The sulfur atoms of the M267 side chains are shown as spheres (enlarged for clarity), with the blue spheres pointing inward while the red spheres are pointing outward relative to the buried binding site. The binding site undergoes an associated conformational change, which is evident from the differences in the surface representation. A movie of the meta-eABF transition is provided in the Supporting Information.

Ideally, a thicker tube would be preferred to generate more sampling along the orthogonal directions to the path and therefore more reasonable free energy estimates. Here, we found that trajectory-based paths could help. We took the trajectory from the 200 ns meta-eABF simulation that used the aforementioned path and analyzed the ≈4,000 frames using the procedure described in Section 2.3. From that we picked a new path, which included all its nodes from a real simulation and utilized it in subsequent meta-eABF simulations. The major benefit we saw in these simulations was that we could relax the tube restraint from 0.2 to 0.8 Å radius (we tried even thicker tubes but they crashed the simulation), and as shown in Figure 10 the PMF barrier between the Met-in and Met-out conformations was lowered markedly using this stage-3 path compared to the barrier computed with the stage-2 path that needed the ultra-thin tube. We note that this procedure is inherently iterative and work is underway to apply further iterations to bring down the PMF to thermodynamically relevant heights.

**Figure 10:**
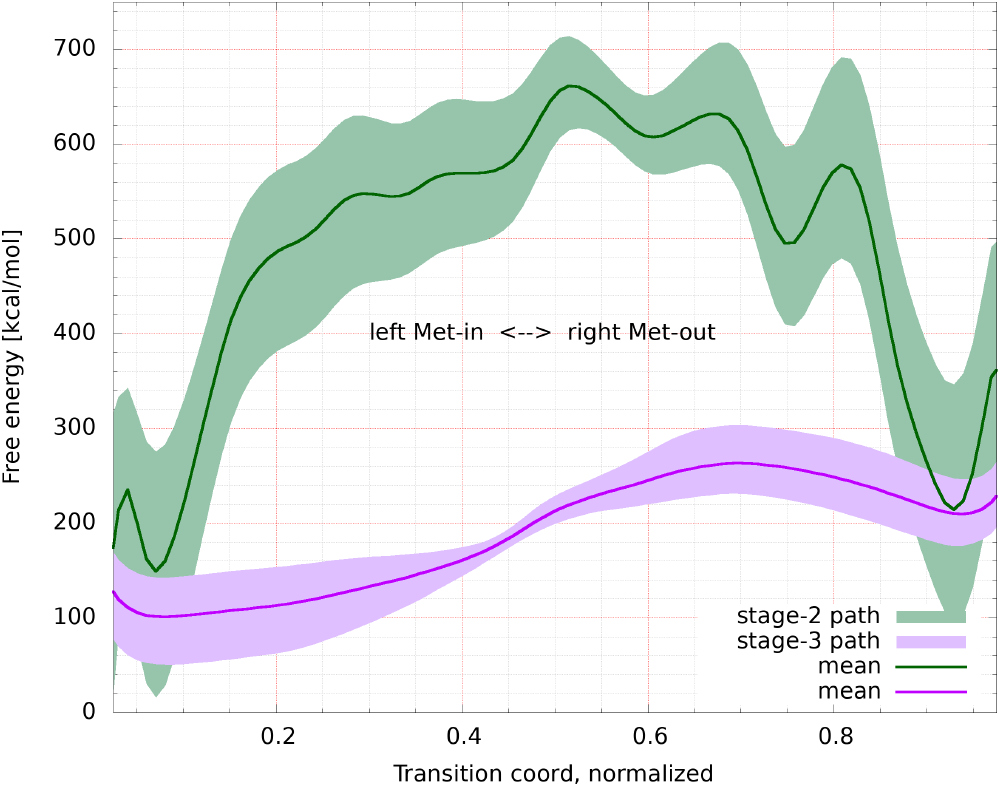
Average PMF of 10,000 boot strapped samples at 95% confidence interval. The horizontal axis represents the transition coordinate of the conformational transformation between the Met-in (left) and Met-out (right) conformations of STING. The vertical axis shows the free energy in kcal/mol. The PMF plot includes the average and the associated 95% confidence interval computed with two different paths. Both paths include the CA atoms and the sulphur atom of Met267, but the stage-2 path is comprised of energy-minimized structures whereas the stage-3 path is constructed from trajectory frames of the very same meta-eABF simulation that employed the stage-2 path.

We conclude with a noteworthy observation that was not clear *a priori*. The STING dimer is symmetrical in both Met-in and Met-out conformations. Therefore, the AA-path algorithm in 2.3 will generate a symmetrical path where the Met267 flip will happen simultaneously in both units. However, there is no physical or biological evidence to assume that the in-out transition should be symmetrical and, in fact, we did observe that the meta-eABF trajectory displayed asynchronous flips as shown in our movie in the SI.^33^ So, besides the trajectory-frame based path being smoother, it can also capture certain aspects of physical reality that cannot be considered by any path method solely based on two endpoint structures.

## 4 Conclusion

Accurate and efficient prediction of conformational changes in biological systems (both the structures and the underlying free energy surfaces) is an important area of research with vast applications in drug discovery. While great strides have been made in protein structure prediction (e.g. AlphaFold, ^42^ RoseTTAFold,^43^ and OpenFold^44^), progress on conformational sampling of biologically-relevant motions has not witnessed a comparable acceleration from artificial intelligence (AI) and machine learning (ML), although some encouraging approaches are beginning to arise. ^45–47^ One reason is that there is much less data associated with protein motion as compared with static structures. One of the few experimental methods that can assess protein motion and energies is NMR,^48^ which is challenging, time consuming, and limited data is available to date. We expect AI and ML to play an increasingly important role in conformational sampling and we anticipate physics-based simulations to play an important role for the foreseeable future. Indeed, physics-based simulations could provide the scale and quality of data necessary for an AI/ML approach.

In this work we presented a new path collective variable (PCV) algorithm built from a smooth ANM (SANM) and includes the option to build an all-atom (AA) path as opposed to a CA-path. We also introduced the concept of refining path CVs iteratively utilizing trajectory frames extracted from a series of high-quality explicit solvent simulations. Furthermore, we pointed out the need for a new kind of restraint for molecular dynamics simulations, which we formulated as a reciprocal barrier restraint. While this type of restraint (we term it ReBaCon for Reciprocal Barrier Constraint) has significant challenges in nonlinear optimization, we showed that it is well-suited for molecular dynamics simulations. In particular, we have found it to be the best approach for path-based enhanced sampling simulations of complex biological conformational changes.

In this work we applied this new PCV and ReBaCon within a hybrid MD sampling method that combines well-tempered metadynamics with extended adaptive biasing force (meta-eABF). We applied our new methods to three complex systems of significant pharmaceutical interest: (1) simulated large-scale conformational changes in the Cullin-RING ligase supramolecular structure responsible for ubiquitination, (2) computationally reproduced the experimental finding that certain small molecules in complex with the oncogenic V617F mutant of the JAK2 pseudokinase domain can allosterically modulate the free energy landscape toward the wild-type conformation, and finally, (3) simulated the conformational change of the STING protein upon activation with particular focus on a specific loop transition. We encourage others in the field to try these methods and propose further improvements, with a focus on biologically relevant problems as opposed to model systems. We are currently using the methods presented here on a variety of conformational sampling problems in our drug discovery programs, including conformational activation and allosteric inhibition.

## Acknowledgement

The authors thank Andra’s Aszódi for help with the CA to BB+CB conversion software and for discussions regarding the statistical error estimates, Fabio Trovato for critical reading of the manuscript and creating the Cullin-RING trajectory movie, and Asghar Razavi for help with the Cullin-RING simulations.

## Supporting Information Available

Movie files of our meta-eABF simulations of the opening/closing of the Cullin-RING complex^32^ (red POI, blue VHL, cyan Elongin, grey Cul2, green Ubiquitin, orange EBE2D2, brown NEDD8, and yellow RBX1), see Section 3.1, and the Met-in to Met-out conformational transition of STING,^33^ see Section 3.3.

## References

(1) Casaletto, J.; Maglic, D.; Toure, B. B.; Taylor, A.; Schoenherr, H.; Hudson, B.; Bruderek, K.; Zhao, S.; O’Hearn, P.; Gerami-Moayed, N. et al. RLY-4008, a novel precision therapy for FGFR2-driven cancers designed to potently and selectively inhibit FGFR2 and FGFR2 resistance mutations. Cancer Research 2021, 81, 1455–1455.

(2) Yan, X.-E.; Ayaz, P.; Zhu, S.-J.; Zhao, P.; Liang, L.; Zhang, C. H.; Wu, Y.-C.; Li, J.-L.; Choi, H. G.; Huang, X. et al. Structural Basis of AZD9291 Selectivity for EGFR T790M. Journal of Medicinal Chemistry 2020, 63, 8502–8511, PMID: 32672461.

(3) Cournia, Z.; Allen, B. K.; Beuming, T.; Pearlman, D. A.; Radak, B. K.; Sherman, W. Rigorous Free Energy Simulations in Virtual Screening. Journal of Chemical Information and Modeling 2020, 60, 4153–4169, PMID: 32539386.

(4) Sittel, F.; Stock, G. Perspective: Identification of collective variables and metastable states of protein dynamics. J. Chem. Phys. 2018, 149, 150901.

(5) Fiorin, G.; Klein, M. L.; H’enin, J. Using collective variables to drive molecular dynamics simulations. Mol. Phys. 2013, 111, 3345–3362.

(6) Elber, R.; Karplus, M. A method for determining reaction paths in large molecules: Application to myoglobin. Chemical Physics Letters 1987, 139, 375–380.

(7) J’onsson, H.; Mills, G.; Jacobsen, K. W. Classical and quantum dynamics in condensed phase simulations; World Scientific, 1998; pp 385–404.

(8) Kirmizialtin, S.; Elber, R. Revisiting and computing reaction coordinates with directional milestoning. The journal of physical chemistry A 2011, 115, 6137–6148.

(9) Templeton, C.; Chen, S.-H.; Fathizadeh, A.; Elber, R. Rock climbing: A local-global algorithm to compute minimum energy and minimum free energy pathways. The Journal of Chemical Physics 2017, 147, 152718.

(10) Das, A.; Gur, M.; Cheng, M. H.; Jo, S.; Bahar, I.; Roux, B. Exploring the conformational transitions of biomolecular systems using a simple two-state anisotropic network model. PLoS Comput. Biol. 2014, 10, e1003521.

(11) Branduardi, D.; Gervasio, F. L.; Parrinello, M. From A to B in free energy space. J. Chem. Phys. 2007, 126, 054103.

(12) Leines, G. D.; Ensing, B. Path finding on high-dimensional free energy landscapes. Phys. Rev. Lett. 2012, 109, 020601.

(13) P’erez de Alba Ortíz, A.; Tiwari, A.; Puthenkalathil, R.; Ensing, B. Advances in enhanced sampling along adaptive paths of collective variables. J. Chem. Phys. 2018, 149, 072320.

(14) Allen, B. K.; Kulkarni, M. M.; Chamberlain, B.; Dwight, T.; Koh, C.; Samant, R.; Jernigan, F.; Rice, J.; Tan, D.; Li, S. et al. Design of a systemic small molecule clinical STING agonist using physics-based simulations and artificial intelligence. bioRxiv 2022,

(15) Andersen, H. C. Rattle: A “velocity” version of the shake algorithm for molecular dynamics calculations. J. Comput. Phys. 1983, 52, 24–34.

(16) Krautler, V.; Van Gunsteren, W. F.; Hünenberger, P. H. A fast SHAKE algorithm to solve distance constraint equations for small molecules in molecular dynamics simulations. J. Comput. Chem. 2001, 22, 501–508.

(17) Bonomi, M.; Branduardi, D.; Bussi, G.; Camilloni, C.; Provasi, D.; Raiteri, P.; Donadio, D.; Marinelli, F.; Pietrucci, F.; Broglia, R. A. et al. PLUMED: A portable plugin for free-energy calculations with molecular dynamics. Comput. Phys. Commun. 2009, 180, 1961–1972.

(18) Tribello, G. A.; Bonomi, M.; Branduardi, D.; Camilloni, C.; Bussi, G. PLUMED 2: New feathers for an old bird. Comput. Phys. Commun. 2014, 185, 604–613.

(19) Bonomi, M. Promoting transparency and reproducibility in enhanced molecular simulations. Nat. Methods 2019, 16, 670–673.

(20) Nocedal, J.; Wright, S. J. Numerical Optimization, 2nd ed.; Springer: New York, NY, USA, 2006.

(21) Lewis, R. M.; Torczon, V. Pattern search methods for linearly constrained minimization. SIAM. J. Optim. 2000, 10, 917–941.

(22) Lewis, R. M.; Shepherd, A.; Torczon, V. Implementing generating set search methods for linearly constrained minimization. SIAM J. Sci. Comput. 2007, 29, 2507–2530.

(23) Eastman, P.; Swails, J.; Chodera, J. D.; McGibbon, R. T.; Zhao, Y.; Beauchamp, K. A.; Wang, L.-P.; Simmonett, A. C.; Harrigan, M. P.; Stern, C. D. et al. OpenMM 7: Rapid development of high performance algorithms for molecular dynamics. PLoS Comput. Biol. 2017, 13, e1005659.

(24) Case, D. A.; Cheatham III, T. E.; Darden, T.; Gohlke, H.; Luo, R.; Merz Jr, K. M.; Onufriev, A.; Simmerling, C.; Wang, B.; Woods, R. J. The Amber biomolecular simulation programs. Journal of computational chemistry 2005, 26, 1668–1688.

(25) Salomon-Ferrer, R.; Case, D. A.; Walker, R. C. An overview of the Amber biomolecular simulation package. Wiley Interdisciplinary Reviews: Computational Molecular Science 2013, 3, 198–210.

(26) Maier, J. A.; Martinez, C.; Kasavajhala, K.; Wickstrom, L.; Hauser, K. E.; Simmerling, C. ff14SB: improving the accuracy of protein side chain and backbone parameters from ff99SB. Journal of chemical theory and computation 2015, 11, 3696–3713.

(27) Wang, J.; Wolf, R. M.; Caldwell, J. W.; Kollman, P. A.; Case, D. A. Development and testing of a general amber force field. Journal of computational chemistry 2004, 25, 1157–1174.

(28) PLUMED-NEST: Reciprocal barrier restraint. Application to path-meta-eABF. https://www.plumed-nest.org/eggs/ 22/032/, accessed by 03-26-2023. PLUMED-NEST: JAK2 2D meta-eABF PMF with statistical analysis.https://www.plumed-nest.org/eggs/23/012/, accessed by 03-26-2023.

(29) PLUMED-NEST: Path meta-eABF simulation of large scale conformational change in STING protein. https://www.plumed-nest.org/eggs/23/013/, accessed by 03-26-2023.

(30) PLUMED-NEST: Path meta-eABF simulation of large scale conformational change in STING protein. https://www.plumed-net.org/eggs/23/013/, accessed by 03-26-2023.

(31) Smooth anisotropic network model (SANM) protein conformational transition path generator. https://githubcom/Psivant/sanm-path-gen., accessed by 03-26-2023.

(32) Movie of the molecular dynamics simulation of the opening and closing of the Cullin-RING protein complex. https://doi.org/10.5281/zenodo.7750767, accessed by 03-26-2023.

(33) Movie of the molecular dynamics simulation of a crucial step in the activation of the STING protein. https://doi.org/10.5281/zenodo.7750819, accessed by 03-26-2023.

(34) Comer, J.; Gumbart, J. C.; H’enin, J.; Lelievre, T.; Pohorille, A.; Chipot, C. The adaptive biasing force method: Everything you always wanted to know but were afraid to ask. J. Phys. Chem. B 2015, 119, 1129–1151.

(35) Lesage, A.; Lelievre, T.; Stoltz, G.; H’enin, J. Smoothed biasing forces yield unbiased free energies with the extended-system adaptive biasing force method. J. Phys. Chem. B 2017, 121, 3676–3685.

(36) Fu, H.; Zhang, H.; Chen, H.; Shao, X.; Chipot, C.; Cai, W. Zooming across the free-energy landscape: shaving barriers, and flooding valleys. J. Phys. Chem. Lett. 2018, 9, 4738–4745.

(37) Fu, H.; Shao, X.; Cai, W.; Chipot, C. Taming rugged free energy landscapes using an average force. Acc. Chem. Res. 2019, 52, 3254–3264.

(38) Dixon, T.; MacPherson, D.; Mostofian, B.; Dauzhenka, T.; Lotz, S.; McGee, D.; Shechter, S.; Shrestha, U. R.; Wiewiora, R.; McDargh, Z. A. et al. Predicting the structural basis of targeted protein degradation by integrating molecular dynamics simulations with structural mass spectrometry. Nature communications 2022, 13, 5884.

(39) McNally, R.; Li, Q.; Li, K.; Dekker, C.; Vangrevelinghe, E.; Jones, M.; Chene, P.; Machauer, R.; Radimerski, T.; Eck, M. J. Discovery and structural characterization of ATP-site ligands for the wild-type and V617F mutant JAK2 pseudokinase domain. ACS Chemical Biology 2019, 14, 587–593.

(40) Drygas, H. On the relationship between the method of least squares and Gram– Schmidt orthogonalization. Acta et Commentationes Universitatis Tartuensis de Mathematica 2011, 15, 3–13.

(41) Steiner’s theorem and least squares. https://people.maths.bris.ac.uk/~mb13434/Steiner_thm.pdf, accessed by 03-26-2023.

(42) Jumper, J.; Evans, R.; Pritzel, A.; Green, T.; Figurnov, M.; Ronneberger, O.; Tunyasuvunakool, K.; Bates, R.; Žídek, A.; Potapenko, A. et al. Highly accurate protein structure prediction with AlphaFold. Nature 2021, 596, 583–589.

(43) Baek, M.; DiMaio, F.; Anishchenko, I.; Dauparas, J.; Ovchinnikov, S.; Lee, G. R.; Wang, J.; Cong, Q.; Kinch, L. N.; Schaeffer, R. D. et al. Accurate prediction of protein structures and interactions using a three-track neural network. Science 2021, 373, 871–876.

(44) OpenFold: A trainable, memory-efficient, and GPU-friendly PyTorch reproduction of DeepMind’s AlphaFold 2. https://zenodo.org/record/6683680#.Yy0kktXMLQN, accessed by 03-26-2023.

(45) Smith, Z.; Ravindra, P.; Wang, Y.; Cooley, R.; Tiwary, P. Discovering Protein Conformational Flexibility through Artificial-Intelligence-Aided Molecular Dynamics. The Journal of Physical Chemistry B 2020, 124, 8221–8229, PMID: 32841026.

(46) Jung, H.; Covino, R.; Hummer, G. Artificial Intelligence Assists Discovery of Reaction Coordinates and Mechanisms from Molecular Dynamics Simulations. 2019,

(47) Martin, W.; Sheynkman, G.; Lightstone, F. C.; Nussinov, R.; Cheng, F. Interpretable artificial intelligence and exascale molecular dynamics simulations to reveal kinetics: Applications to Alzheimer’s disease. Current Opinion in Structural Biology 2022, 72, 103–113.

(48) Jiang, Y.; Kalodimos, C. G. Molecular Chaperones: Confirmation for conformational selection. eLife 2018, 7, e34923.

